# Integration of Deep-Learning and Species Distribution Models for Classification of Animal Species of the Brazilian Fauna

**DOI:** 10.64898/2026.05.06.723365

**Authors:** Mateus Braga Oliveira, Heder Soares Bernardino, Alex Borges Vieira, Antônio Arbex Barroso, Douglas A. Augusto

**Affiliations:** Universidade Federal de Juiz de Fora, Juiz de Fora, MG, Brazil; Oswaldo Cruz Foundation, Av. Brasil, 4365, Manguinhos, Rio de Janeiro, Brazil

**Keywords:** Brazilian fauna, animal classification, citizen science, convolutional neural network, species distribution model, genetic algorithm

## Abstract

The automated classification of animals from photos is important in ecology and conservation biology for organizing and understanding the immense diversity of species, as well as facilitating effective conservation and management practices. It is equally important for disease surveillance systems, allowing prompt detection of anomalies in species distributions and boosting citizen-scientist platforms by making user-reported data more accurate and convenient. Image classification uses photos and can also rely on the geographical locations of animals to improve performance. While image classification models have difficulties in classifying low-quality images, unbalanced datasets, and with a small number of images, species distribution models have difficulty in classifying species that coexist in a given region. We propose here strategies for combining image classification models based on deep neural networks with species distribution models using genetic algorithms. The proposal is applied to a real-world dataset comprising fifteen classes of animals from the Brazilian fauna obtained from Fiocruz’s citizen-scientist Wildlife Health Information System (SISS-Geo). The SISS-Geo photos portray the reality of animals in their environments, with varying quality, and pose numerous difficulties for classification. Experimental results demonstrate that the proposed integration consistently outperforms standalone models. While individual SDMs achieve Top-1 accuracies of 27.79% (MaxEnt) and 31.76% (Bioclim), and CNN-based classifiers reach 58.17% with ResNet50 and 64.13% with ResNet-152, the hybrid strategies yield substantial improvements. The genetic algorithm–based integration with a single global weight achieves up to 67.96% Top-1 accuracy, whereas the class-specific integration using fifteen parameters attains the best overall performance, reaching 69.03%.

## 1 INTRODUCTION

Two fundamental aspects of environmental conservation are species classification and species distribution, which focus on recognizing different animal species and understanding the geographic areas where they are found [1]. With this information, it is possible to determine the existence of endangered and invasive species, as well as to evaluate the potential reintroduction of species in environments affected by natural disasters or environmental tragedies. Another important component is the continuous monitoring of wildlife to identify possible diseases, especially those that can infect humans (zoonoses), such as sylvatic yellow fever. This enables early detection and effective control of disease outbreaks, helping to protect both human and animal lives.

Technological advances in recent years have made several applications feasible. The use of deep neural networks for image classification has become a widespread and effective approach [2, 3]. In parallel, wildlife monitoring programs increasingly employ camera traps in forests and protected areas, generating large volumes of animal images in their natural habitats. Moreover, advances in mobile technology and its ubiquity have enabled citizens to increasingly record wildlife using smartphones and cameras in both urban and rural environments.

In addition to image data, a substantial amount of information is available regarding the geographic locations of wildlife records. Many observations are associated with precise geographic coordinates, creating datasets that combine visual information with spatial occurrence data. Together with increasingly accurate environmental variables provided by environmental agencies, this has driven the development of Species Distribution Modeling (SDM) and Ecological Niche Modeling (ENM) [4]. These models use environmental variables and species occurrence data to estimate the potential distribution of species across landscapes, enabling a wide range of ecological applications, including the assessment of environmental effects on species presence, the detection of invasive species, and the analysis of migration and range shifts [5].

Although deep learning-based image classification is widely considered the most accurate approach for species identification from images, species distribution models can outperform purely visual classifiers in challenging scenarios, particularly when image quality is low or when morphologically similar species are involved [6]. Both approaches are therefore complementary. However, their combined use remains relatively uncommon in the literature [1]. Existing works [6, 7, 8, 9] have explored multimodal approaches using image and georeferenced data, but typically focus on restricted geographic regions, a small number of visually similar species, and often rely on heavily curated datasets with low-quality images removed.

Here, we present an integrated framework for animal species classification that combines deep learning and species distribution modeling, focusing on a challenging, real-world, citizen-science dataset from a wildlife health surveillance platform characterized by high variability in image quality. We employ two deep convolutional neural networks, ResNet-50 and ResNet-152, for image-based species classification, and two SDM approaches, MaxEnt and Bioclim, for modeling species distributions.

The integration of models is performed using a genetic algorithm, which combines parameters that control the relative contribution of each model [10]. This evolutionary optimization process searches for the weight configurations that maximize classification performance on validation data, enabling a data-driven and adaptive fusion of visual and environmental information. Two different strategies are adopted for combining the models using a genetic algorithm. In the first strategy, the combination is controlled by a single global parameter, which is applied uniformly across all classes. In the second strategy, the genetic algorithm is used to estimate 15 distinct parameters for the model combination, with one specific parameter assigned to each class, allowing a more flexible and class-dependent weighting of the models.

We evaluate the proposed framework using data from SISS-Geo, a project based on real-world records of Brazilian fauna that incorporates a citizen science approach, engaging the public in biodiversity data collection [11]. We then assess the framework using widely adopted metrics in the literature, including Top-1 accuracy, precision, and Top-N accuracy. The results are presented through comparative analyses among individual models and the integrated approach, highlighting performance gains, trade-offs, and generalization behavior across species and regions.

Thus, the main contributions of this work are: (1) the use of real-world data from an official wildlife health surveillance platform in Brazil, incorporating records of Brazilian fauna with a strong emphasis on zoonotic disease monitoring; (2) the proposal of an integrated multimodal framework that combines deep convolutional neural networks with species distribution models; and (3) the application of genetic algorithms to optimize the combination of heterogeneous models, enabling flexible and adaptive integration strategies; and (4) the demonstration of the socio-environmental relevance of automated species identification and distribution modeling, supporting biodiversity conservation, wildlife health surveillance, and decision-making processes related to zoonotic disease prevention and environmental management.

Automatic classification of wildlife records is essential for monitoring species, assessing their health conditions, detecting changes in species distributions, and strengthening biodiversity conservation efforts. Furthermore, it enables more efficient and accurate data collection by non-specialists, which is critical for the scalability and reliability of citizen-science platforms. Finally, these advances also contribute to public health and to best practices in sustainable development.

## 2 MATERIALS AND METHODS

This section describes the materials, data sources, and methodological components required to develop the proposed integrated framework. We combine deep-learning–based image classification models with species distribution models to address a challenging real-world problem involving wildlife records in Brazil. The approach leverages animal photographs, georeferenced occurrence data, and environmental information to jointly identify species and model their spatial distribution. Such integration is particularly relevant in biodiversity-rich regions, where correct species identification and distribution mapping are essential for conservation planning, ecological monitoring, and wildlife health surveillance.

The experiments are conducted using data from the SISS-Geo platform, a nationwide citizen-science system that collects wildlife images, geographic coordinates, temporal metadata, and contextual information related to animal health and environment. The dataset reflects realistic conditions, including heterogeneous image quality, class imbalance, and the coexistence of multiple species within the same regions. These characteristics make the task especially challenging and distinguish it from more controlled benchmarks commonly used in the literature. The section therefore, outlines the data characteristics, the selected image classification and species distribution models, and the strategy adopted to integrate their outputs into a unified framework.

### 2.1 Dataset

SISS-Geo is a collaborative platform for wildlife monitoring [12] led by Fiocruz whose main goal is to produce early alerts of emerging diseases so that health authorities can act to prevent or mitigate their impact on humans, animals, and on the environment. Citizen scientists volunteers and health professionals use SISS-Geo to register occurrences involving animals, their conditions, their precise location, and information regarding the surroundings. Records can be made by the SISS-Geo^1^ mobile applications (Android or iOS), for convenience in fieldwork, or through a Web platform that also offers many analysis and exportation tools.

Once records are sent to SISS-Geo, predefined alert rules and computational models predict the seriousness of the situation. Examples of alarming situations include the observation of one or more dead monkeys or multiple sick animals found in a given region in a short period of time.

If an alert is issued, health authorities are immediately informed and, depending on their judgment, they go on-site to check the situation and, in some cases, collect animal samples for laboratory analysis — the resulting diagnostics feed back into SISS-Geo to train the alert models.

Figure 1 depicts the overall workflow of SISS-Geo, including the input of records, alert system, laboratory diagnostics, and model training.

**FIGURE 1.**
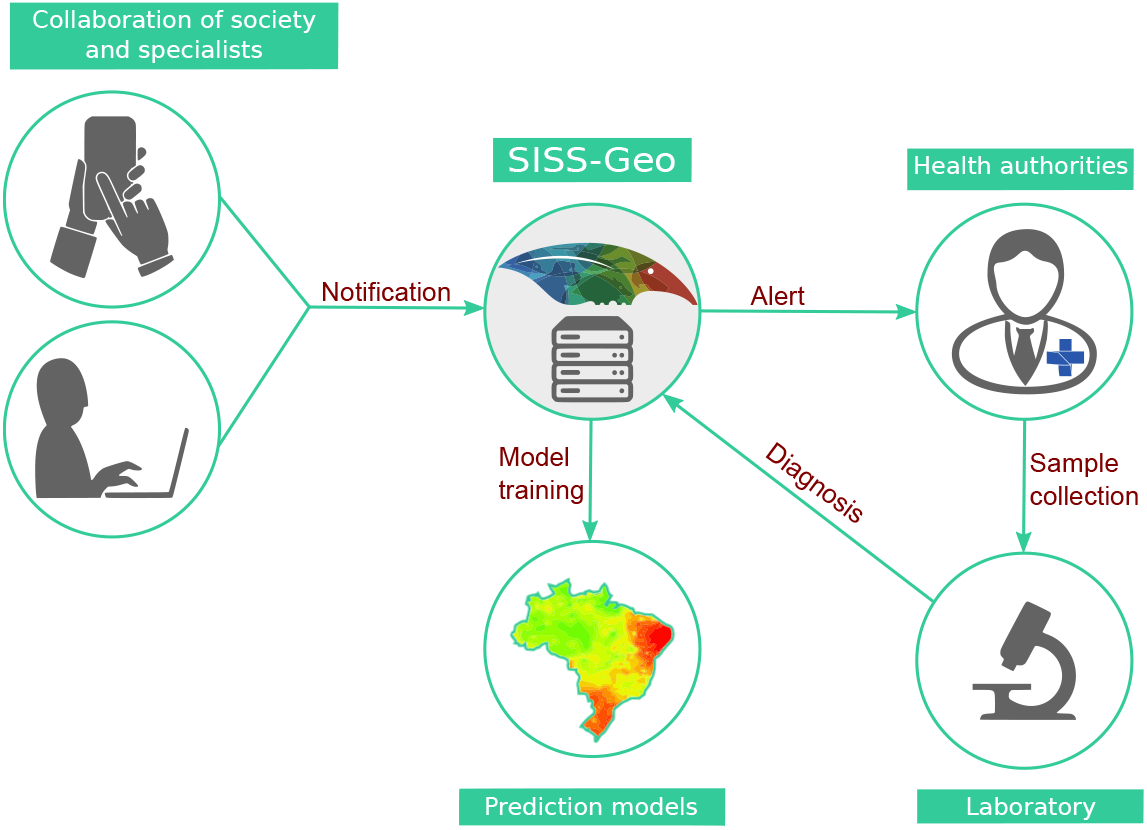
SISS-Geo’s overall workflow.

A key aspect of SISS-Geo that sets it apart from related projects (such as iNaturalist [13]) is the fact that it is embraced by governmental and non-governmental health institutions and aims at supporting decision-making regarding human and animal health. The platform features dedicated modules for health authorities, such as alert notification and disease input from diagnostics. From the produced data, prediction models of zoonoses occurrence are trained and applied to estimate the suitability of regions for diseases so that, for instance, vaccination campaigns can be planned in advance.

In SISS-Geo, almost all animal records are from the Brazilian fauna. Although the majority of the records are curated by specialists and have assigned the animal’s species, here we deal with animal types, with each type representing a group of related species. In addition to the photos of the animals, SISS-Geo stores other relevant data, notably the location where the record was made and detailed characteristics of the surrounding region. In this work, besides the animal type (class), only the images and location of the registered animals are used.

The records used here were obtained from the general dataset of SISS-Geo, comprising 15 classes (out of a total of 40 classes). We selected these classes due to their availability of data. In Figure 2, the distribution of records among the working classes is presented. From this figure, one can verify that the data is highly unbalanced, a common issue when solving real-world problems.

**FIGURE 2.**
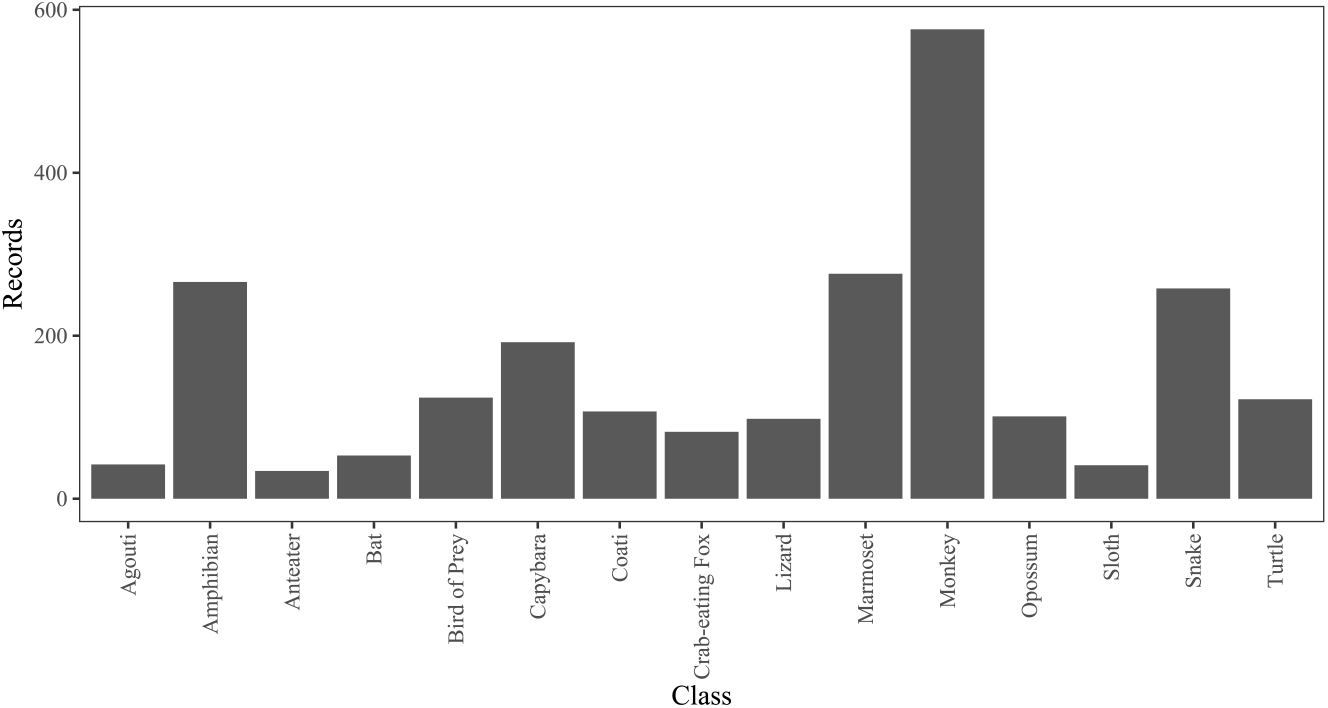
Number of records for each class.

As one can see in Figure 3, the photos depict different environments and shading, in addition to the large distance to the animal registered in the image. It is also possible to see in this figure how the environment is challenging for the correct classification of images. Some animals are disguised in their environment. Other animals registered in SISS-Geo are dead or injured. This is also a challenging factor for the proper classification.

**FIGURE 3.**
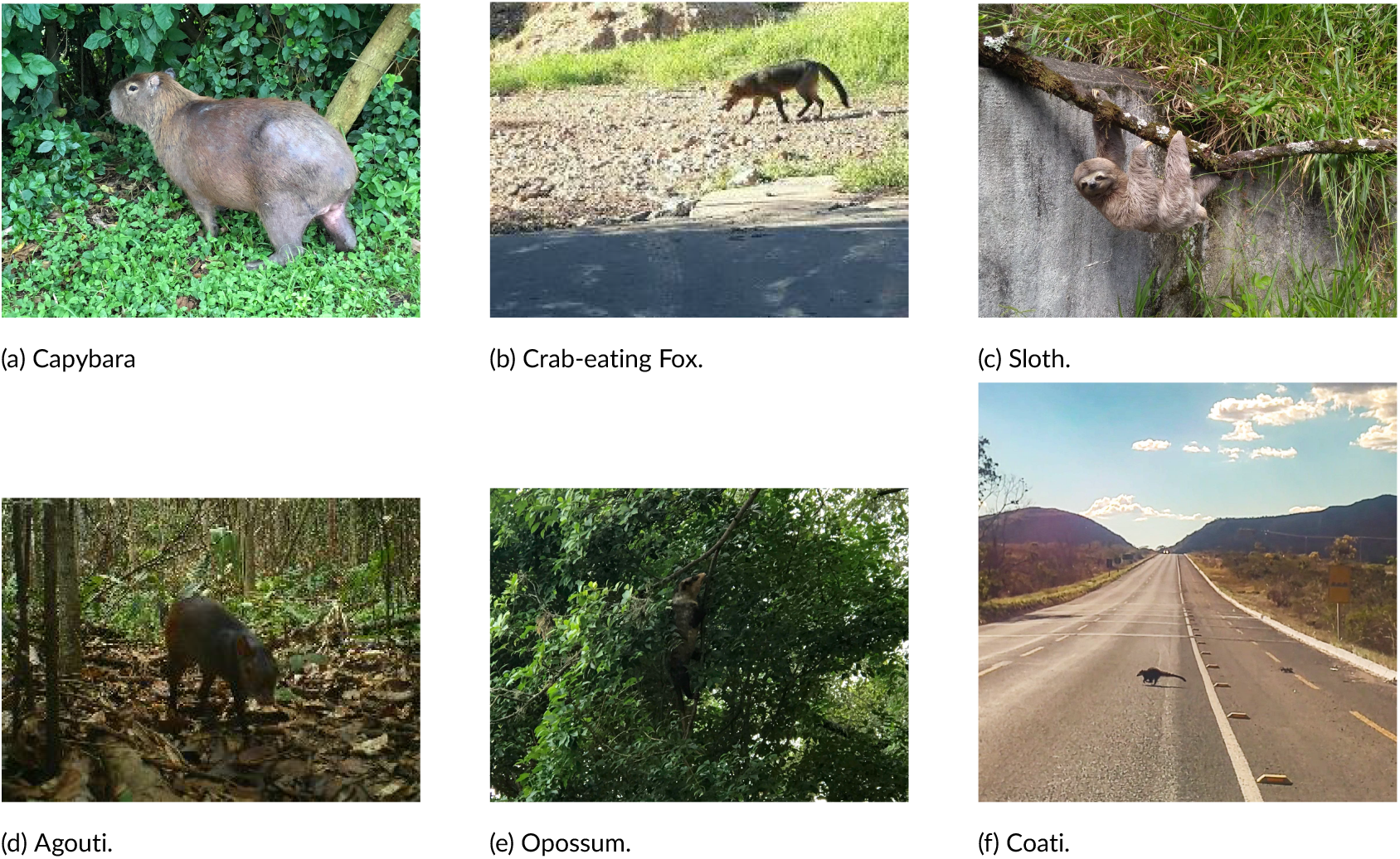
Examples of Animal Classes. Images courtesy of SISS-Geo (user-contributed observations).

For instance, one can see in Figure 4 examples of photos in which the class identification is hard to even for humans. Another aspect when working with data obtained from a citizen-science project — in particular where the object of interest can move — is the diversity in the quality of the images.

**FIGURE 4.**
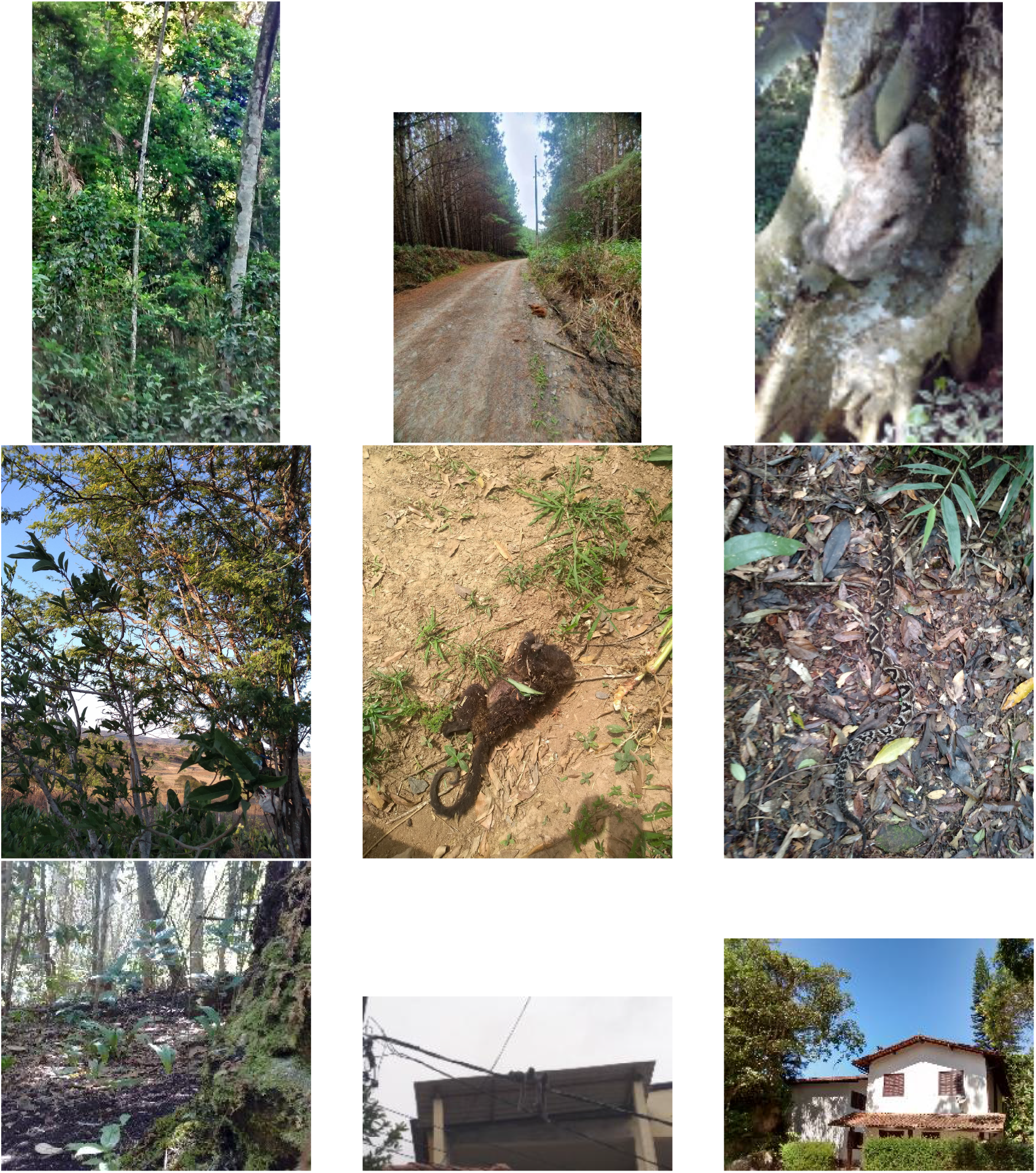
Examples of images that are hard to classify. Images courtesy of SISS-Geo (user-contributed observations).

The difficulty in classification is due to different factors. Among these factors, how far the animal is from the photographer is critical. Often, citizens are not close to the animal they take the photo of, which is mainly a result of local inaccessibility or the fact that the animals tend to run away from people. Another factor that is observed is the condition of the environment, which directly affects the lighting of the image. A third common factor in the SISS-Geo images is the presence of records of dead, decaying animals; they are common as SISS-Geo focuses on sick and dead animals due to their importance as sentinels for the circulation of diseases.

### 2.2 Data Augmentation

To increase the number of images of the SISS-Geo dataset, some processing for data augmentation is carried out. These approaches are: (1) horizontal mirror; (2) random rotation; (3) random zoom; (4) random brightness; and (5) random noise.

To compare the different approaches, a detailed organization of the datasets used here was necessary. In computer vision works, a standard division of the dataset into training, validation, and test sets is usually performed in a random way [14]. However, for an effective comparison between the image classification and species distribution models and the integrated model, it is necessary to ensure that each record used in the test set is composed of an image associated with its geographic location.

Figure 5 illustrates, in an exemplary manner, the standard data augmentation operations applied to an image from SISS-Geo, including transformations such as rotations, reflections, cropping, and geometric and radiometric variations. These operations aim to artificially expand the training dataset while preserving the semantic content of the original scene, thereby increasing variability and improving the robustness and generalization capability of the deep learning models employed.

**FIGURE 5.**
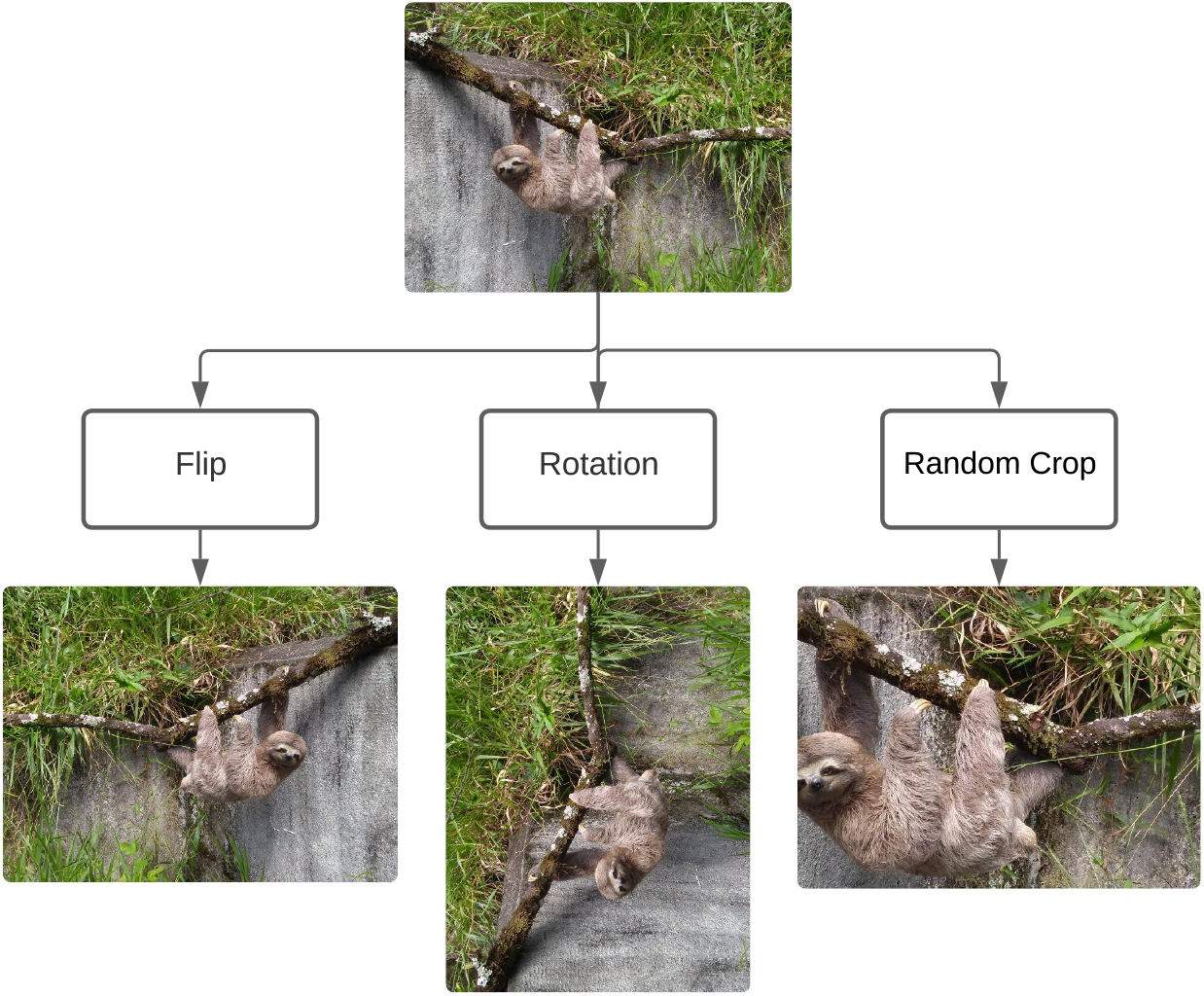
Data Augmentation Operations.

**FIGURE 6.**
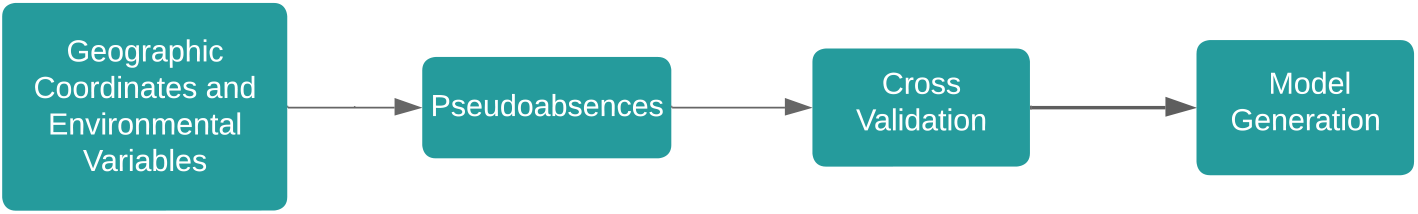
Species Distribution ModelWorkflow.

**FIGURE 7.**
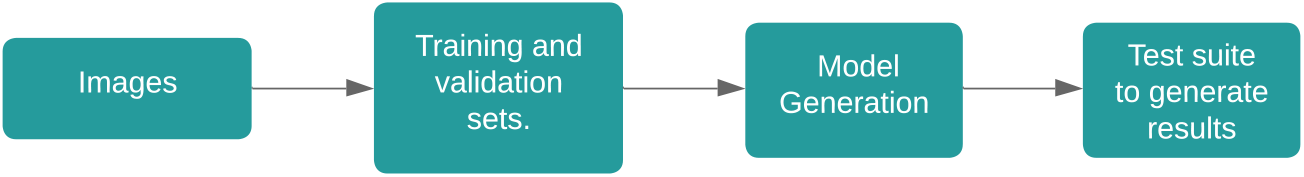
Image Classification ModelWorkflow.

### 2.3 Spatial Distribution of Animal Species

Knowing the geographical potential distribution of species is essential to support evolutionary and ecological studies, including the prediction of occurrences of diseases. However, the lack of data about the actual distribution of species and their unequal spatial concentration makes it difficult to rely on this information for analyses and decision-making. Unfortunately, collecting more data involves laborious and expensive fieldwork and would require constant updates to account for temporal variations.

With this problem in mind, Species Distribution Modeling (SDM) methods have emerged as a great contribution to studies carried out in these areas. This methodology can be applied in different areas. For example, to prioritize areas for conservation [15], discuss biogeographical patterns [16] and, with the availability of modeled data on past and future climate, predict changes in the distribution of organisms over time [17].

Species distribution models (SDMs) aim to identify the environmental conditions necessary for sustaining populations of a target species by establishing associations between environmental variables and species occurrence records. These associations can be determined using various algorithms. SDMs serve multiple purposes, including: 1) Assessing the relative suitability of currently occupied habitats by the species; 2) Estimating the relative suitability of geographic areas not currently occupied by the species, which is relevant for identifying potential distributions or invasive species; 3) Predicting changes in habitat suitability over time under specific scenarios of environmental change; and 4) providing an estimation of the ecological niche of the species.

The applications of SDMs are diverse and involve constructing models that approximate the species’ distribution niche. These models are based on the theoretical framework of the ecological niche, a fundamental concept in ecology and evolutionary biology.

There are two types of biotic data associated with Species Distribution Models: presence records and absence records for a given species. Absence records are extremely rare and were one of the first problems faced by species distribution modeling [18]. Absence data do not always reflect a real absence or unsuitability of the environment for the occurrence of the species, and may simply indicate that the species did not reach a certain location or that there is a lack of inventories in the location. Normally, SDMs are built with only presence data, which can generate other problems during modeling.

The Abiotic Data are the environmental variables. Continuous environmental data are the most used in SDM, such as climate data (temperature and precipitation) and topographic data (elevation and slope of the land).

#### 2.3.1 Environmental Data

WorldClim is a set of global climate layers (gridded climate data in GeoTIFF format) that can be used for mapping and spatial modeling. WorldClim^2^ version 2 [19] contains gridded monthly average climate data for the period 1970-2000 with different spatial resolutions, from 30 seconds (about 1 *k m*^2^) to 10 minutes (about 340 *k m*^2^).

The dataset includes the main climatic variables (monthly minimum, average and maximum temperature, precipitation, solar radiation, wind speed, and water vapor pressure) as well as 19 derived bioclimatic variables (annual average temperature, average diurnal interval, isothermality, temperature seasonality, maximum temperature, temperature of the hottest month, minimum temperature of the coldest month, annual temperature range, average temperature of the wettest quarter, average temperature of the driest quarter, average temperature of the warmest quarter, average temperature of the coldest quarter, annual precipitation, precipitation of the wettest month, precipitation of the dry month, precipitation seasonality (coefficient of variation), precipitation of the wettest quarter, precipitation of the driest quarter, precipitation of the hottest quarter, precipitation of the warmest quarter cold).

In this work, we use WorldClim2 bioclimatic environmental variables. We chose to use WorldClim because the 19 bioclimatic variables are a popular choice for bioclimatic SDMs in the specialized literature.

#### 2.3.2 Species Distribution Models

Species distribution models [20] are mainly based on environmental conditions and are generated from a set of rules ranging from simpler mathematical solutions (Euclidean Distance, Bioclim), to statistical adjustments (Generalized Linear Models – GLM, Models Generalized Additives - GAM) to algorithms derived from artificial intelligence and search (Maxent, GARP, Neural Networks). What these algorithms calculate is the environmental similarity between the known places of occurrence for the species and other regions still unknown. In the end, the places with the greatest similarity are considered areas with a high probability of occurrence.

Maximum Entropy - MaxEnt, uses the principle of maximum entropy in presence data to estimate a set of functions that relate to environmental variables of the habitat to approximate the potential geographic distribution of the species [21]. The principle of maximum entropy, which says that the best approximation to an unknown probability distribution is the one that satisfies any restriction on the distribution.

One of the steps to improve the quality of MaxEnt training is the cross-validation process. Cross-validation is a widely used technique for evaluating the performance of machine learning models and consists of partitioning the data into sets (parts), where one set is used for testing to evaluate the performance of the model, while the remaining sets are used for training the model.

In the present work, a buffer-spacing strategy was used at each georeferenced point used in the MaxEnt models to limit data density, thus mitigating the excessive influence of similar points while preserving their diversity. This methodology was applied to standardize the number of records available for use in species distribution models.

By generating pseudoabsence data as part of its workflow, MaxEnt has the advantage of being capable of using presence-only data, therefore not requiring confirmed absence data from specific areas. The generation of pseudoabsence employed was based on the environmental distance between the presence records, considering regions where the climatic condition is favorable for the existence or absence of the class under analysis. We set the number of generated pseudoabsence data to 400 points for each class.

The Bioclim model is one of the earliest and most widely known approaches in species distribution modeling (SDM), also referred to as a *climate-envelope model*. Originally developed at the Commonwealth Scientific and Industrial Research Organisation (CSIRO^3^) in Australia during the 1980s [22], Bioclim uses only species occurrence data (presence-only) and a set of continuous environmental variables to define an *n*-dimensional environmental envelope that represents the species’ realized climatic niche.

Bioclim constructs a multidimensional space based on the environmental conditions (e.g., temperature, precipitation) observed at the species’ known occurrence points. For each environmental variable, the algorithm determines threshold values (typically percentiles such as the 5th and 95th) that encompass most occurrences, excluding extreme outliers. Any geographic location whose environmental values fall within this envelope is considered climatically suitable for the species. The degree of suitability can be further quantified by measuring how close a location’s environmental values are to the median values of the known occurrences.

The model is conceptually simple, easy to interpret, and does not require absence data, making it useful for exploratory studies and educational purposes. Species distribution models were developed using the R package dismo^4^. Bioclim is particularly suitable when only presence data and continuous climatic variables are available. It provides a straightforward way to visualize the climatic range within which a species occurs.

Because it assumes all locations within the envelope are equally suitable, Bioclim tends to overestimate potential distributions. It also does not account for interactions among environmental variables, dispersal barriers, biotic inter-actions, or temporal dynamics [22]. Furthermore, the model is limited to continuous variables and performs poorly when extrapolating beyond the observed environmental range, such as in climate change projections.

### 2.4 Image Classification

Image classification [23] is the task of assigning a label or class to an entire image. Image classification models take an image as input and return a prediction of its class, typically also returning the prediction probability for all classes.

Over the years, various convolutional neural network (CNN) architectures have been developed, resulting in significant advancements in the field of deep learning. Several notable CNN architectures, including LeNet-5, AlexNet, VGG, and ResNet, have emerged as leading approaches that have substantially reduced error rates compared to earlier models [24].

However, to obtain high classification accuracy of images, a large amount of data is required. For this, many models that work with Computer Vision problems apply techniques for data augmentation. Thus, this section presents the neural network used in this research, as well as the steps to increase the data.

#### 2.4.1 Residual Neural Network

ResNet, a well-known artificial neural network, is a residual neural network introduced by He et al. in their paper [25]. The powerful representation capabilities of ResNet have been leveraged in numerous computer vision applications. Beyond image classification, ResNet has also proved beneficial for tasks such as face recognition and object detection, showcasing its pioneering contributions in the field.

ResNet-50 is a deep convolutional neural network (CNN) with 50 layers, built on the concept of residual blocks. Residual networks address the challenges of deep networks by incorporating skipping connections, allowing information to flow directly between layers. This approach improves model accuracy and is a key strength of ResNet.

The architecture known as *ResNet-152* is a version of the residual network family introduced by He, Zhang, Ren & Sun [25]. ResNet-152 uses 152 layers, employing “skip” or identity connections so that layers learn the residual function which eases optimization of very deep networks.

Table 1 presents a comparative overview of the ResNet-50 and ResNet-152 architectures, two widely used deep convolutional neural networks based on residual learning. Although both models employ bottleneck residual blocks, they differ significantly in depth, computational cost, and performance.

**TABLE 1.** Comparison between ResNet-50 and ResNet-152.

| Characteristic | ResNet-50 | ResNet-152 |
| --- | --- | --- |
| Depth (layers) | 50 | 152 |
| Architecture type | Bottleneck residual blocks | Bottleneck residual blocks |
| Number of blocks | 16 bottleneck blocks | 50 bottleneck blocks |
| Parameter count | ~25.6M | ~60.2M |
| Top-1 accuracy (ImageNet) | ~76% | ~78.5% |
| Top-5 accuracy (ImageNet) | ~93% | ~94.2% |
| Computational cost (FLOPs) | ~4.1B | ~11.3B |
| Training time | Faster | Slower |
| Inference speed | Faster | Slower |
| Memory usage | Lower | Higher |
| Typical use cases | Real-time tasks, embedded systems | High-accuracy offline tasks |

ResNet-50, with 50 layers and approximately 25.6 million parameters, offers a balanced trade-off between accuracy and efficiency. Its lower computational cost and memory usage make it suitable for real-time applications, embedded systems, and scenarios in which inference speed is a priority. In contrast, ResNet-152 extends the architecture to 152 layers, resulting in more than 60 million parameters and substantially higher computational demands. This increased depth typically leads to improved feature representation and better top-1 and top-5 accuracy on large-scale datasets such as ImageNet.

However, the gains in performance come at the expense of higher training time, slower inference, and greater memory requirements. As a result, ResNet-152 is more commonly applied in offline or high-accuracy tasks where computational resources are less constrained. Overall, the comparison highlights the trade-offs between efficiency and accuracy, guiding the selection of the appropriate architecture based on the specific requirements of a given application.

### 5.2 Genetic Algorithm (GA)

Optimization is a fundamental area in science and engineering that focuses on finding the best possible solution to a problem under a given set of constraints. In many real-world applications, the search space is large, complex, and non-linear, making classical deterministic optimization methods impractical or inefficient. In such contexts, metaheuristic approaches are often adopted to provide approximate yet effective solutions within a reasonable computational time.

Evolutionary computation is a family of population-based optimization techniques inspired by natural evolution and biological processes. These methods, which include genetic algorithms, genetic programming, and evolutionary strategies, operate through mechanisms such as selection, reproduction, mutation, and survival of the fittest. By iteratively evolving a population of candidate solutions, evolutionary algorithms are particularly well-suited for solving complex optimization problems with multiple local optima, noisy objective functions, or heterogeneous solution spaces.

In a genetic algorithm, the process starts with an initial population of chromosomes, which are possible solutions to a given problem. Those chromosomes consist of an array of genes.

The whole optimization problem is encoded into a fitness function, which receives a chromosome and returns a number that tells the fitness (or goodness) of the solution. The higher the fitness, the better the solution encoded in the chromosome.

Then starts the loop. At each iteration (generation), a number of good chromosomes are selected for breeding (parent selection). Parents are combined two-by-two (crossover) to generate new chromosomes (children). The children are finally mutated by modifying somewhat randomly part of their genes, allowing for completely new solutions to emerge. The children go to the next generation, and a new iteration starts.

The genetic algorithm begins with the initialization of a population of randomly generated weights, each constrained to the interval [0, 1]. These weights represent the relative contribution of Model 1 when combining its predicted probabilities with those of Model 2. As in classical evolutionary methods, population size directly affects the diversity of the search space and therefore influences convergence behavior [26]. After initialization, each individual is evaluated by combining model outputs according to its weight, and the resulting accuracy—computed relative to the ground-truth labels—serves as the fitness function. Fitness-driven evaluation is a core component of evolutionary search, enabling the algorithm to progressively focus on more promising regions of the solution space [27].

Once fitness scores are computed, individuals are randomly subjected to a selection process based on pairwise tournaments. In each tournament, the individual with the higher fitness is selected to compose the parental set. This strategy allows individuals with relatively low fitness to be selected when competing against even less fit opponents, therefore preserving genetic diversity within the population throughout the future generations, avoiding premature convergence. New individuals are then generated using recombination of the parental set of individuals, in which pairs of candidates produce offspring whose weights result from averaging parental values. This mechanism allows offspring to inherit characteristics from both parents, consistent with the principles of genetic recombination in evolutionary computation [28]. To prevent stagnation and enhance exploration, mutation introduces small random perturbations to some weights with low probability, an essential mechanism for maintaining genetic diversity within the evolving population. At last, the least fit individuals in the original population are replaced by the best performing offspring. Elite preservation ensures that the best solutions found so far are retained across generations, reducing the risk of regression in performance.

This iterative cycle of evaluation, selection, recombination, and mutation continues for a predetermined number of generations or until a specified stopping criterion is satisfied—such as achieving a target accuracy or observing convergence. At the end of the evolutionary process, the best-performing weight is selected as the optimal solution. This weight is then used to combine the outputs of the two models, enabling evaluation of the ensemble’s performance relative to each model individually. The use of learned weighting strategies in model combination aligns with established ensemble learning principles, where weighted voting and probability combination frequently yield superior predictive performance [29].

#### Algorithm 1: Genetic algorithm used to learn the optimal ensemble weight *w* ∈ [0, 1].

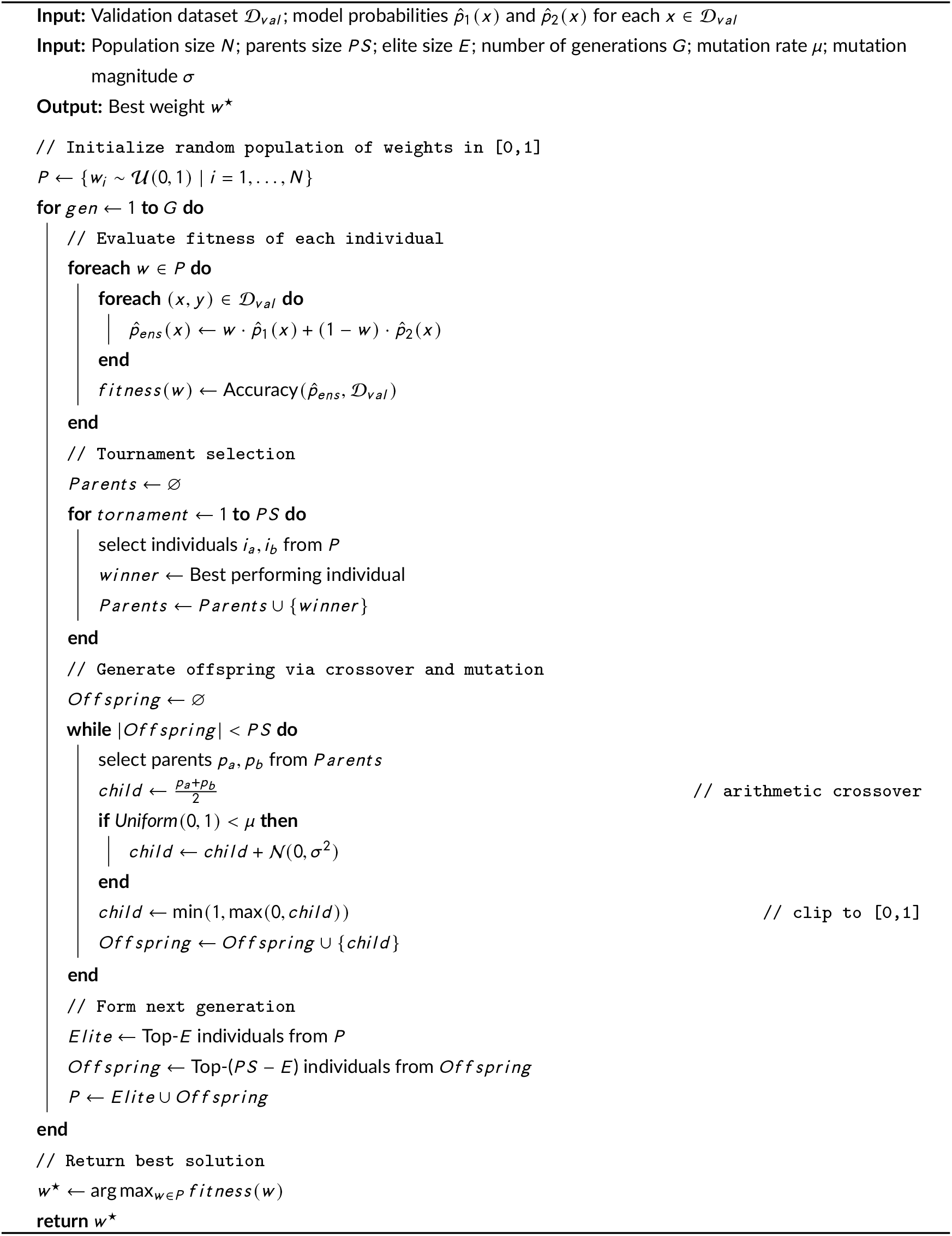

## 3 RELATED WORKS

Some authors only use image location as additional information in the image classification model. In [30], for example, the author does not work with a species distribution model, using geographic coordinates together with images of the species under study. Thus, this work does not take into account the influence that the environment has on the distribution of species. The same was performed by Chu et al. [7], who also use only geographic coordinates to help classify images, using datasets that include animals, plants, and their respective locations.

Terry et al. [6] also cropped their images, centering the animal in the image and resizing the images to a standard format (299 × 299 pixels), in addition to excluding a number of images that could be used in modeling, such as low-quality images.

What can be seen is that, in the literature, images are preprocessed to reduce possible problems in image classification and species distribution models. Together with the preprocessing of the images, the species considered are restricted to a class (such as ladybugs in [6] and Chinese Galliformes in [9]).

SISS-Geo is a single source of data for both the image classification model and the species distribution model because it contains animal records with images and geographic coordinates of their location. Lin et al. [9] used a compilation of data from different sources. It is also possible to aggregate more data from other sources for use in conjunction with SISS-Geo. However, there are possibilities of inserting duplicate or incomplete data. Therefore, when using a single data source, possible errors that can be inserted in the classification are avoided.

Another point that differentiates our work is the fact that Lin et al. use a single group, the Chinese Galliformes. In our work, we focused on classifying different groups of animals from the Brazilian fauna, with different characteristics, that coexist in the same study region, sharing the environment.

Environmental coexistence is another point that stands out in our research. It can be seen in the work by Lin et al. that most of the studied species are endemic, which favors species distribution models to correctly classify each of these groups of animals. It is worth noticing that the NicheNet model by Lin et al. [9] made several misclassifications when classifying species that shared the same environmental region.

In contrast to previous studies, our methodology does not rely solely on the direct incorporation of geographic coordinates as auxiliary features, nor does it focus on a restricted taxonomic group. Instead, we combine image-based classification and species distribution modeling as independent but complementary sources of information. The outputs of these heterogeneous models are subsequently integrated through optimization strategies based on genetic algorithms, allowing the contribution of each model to be adaptively adjusted according to the characteristics of the species being classified. This approach enables the exploitation of both visual and ecological information while addressing the challenges posed by species that coexist in the same environments, resulting in a more flexible and generalizable framework for wildlife species identification.

## 4 METHODOLOGY

This study proposes an integrated modeling framework that combines deep convolutional neural networks for image classification with classical species distribution models and a genetic optimization stage. The overall objective is to merge visual information derived from field images with environmental and climatic descriptors, producing hybrid predictive models that are both accurate and ecologically interpretable.

The methodology accounts for the heterogeneous structure of the dataset, in which each record in the SISS-Geo system may contain either a single image or multiple associated images. Records with multiple images provide natural data diversity, while single-image records are enhanced through an extensive data augmentation pipeline. This ensures that all records contribute comparably to the learning process, preventing bias toward data-rich entries. After balancing the number of effective images per record, the subsequent modeling steps integrate both image-based and environmental information.

The proposed pipeline operates in three main stages. First, ResNet-based convolutional neural networks (ResNet-50 and ResNet-152) are trained to classify the images and generate probabilistic outputs for each class. In parallel, species distribution models (MaxEnt and BioClim) estimate the likelihood of each record belonging to a given class based on environmental parameters. Finally, a genetic algorithm combines the outputs of these two modeling streams to form hybrid models — one using a single global parameter and another using fifteen class-specific parameters — optimizing the integration between visual and environmental predictions.

### 4.1 Data Augmentation of SISS-Geo

To address the inherent imbalance of the SISS-Geo dataset, data augmentation plays a central role in the methodological pipeline. As illustrated in Figure 8, the SISS-Geo recording process generates heterogeneous records in terms of both the number of images per record and their associated geographic information, which naturally leads to class imbalance. In this context, data augmentation is introduced as a systematic strategy to mitigate these disparities by artificially increasing the number of training samples for underrepresented classes, while preserving the semantic and spatial characteristics of the original data. The following subsection details the data augmentation procedures adopted in this work, describing the types of transformations applied and their role in promoting a more balanced dataset, thereby enhancing the robustness and generalization capability of the image classification and species distribution models.

**FIGURE 8.**
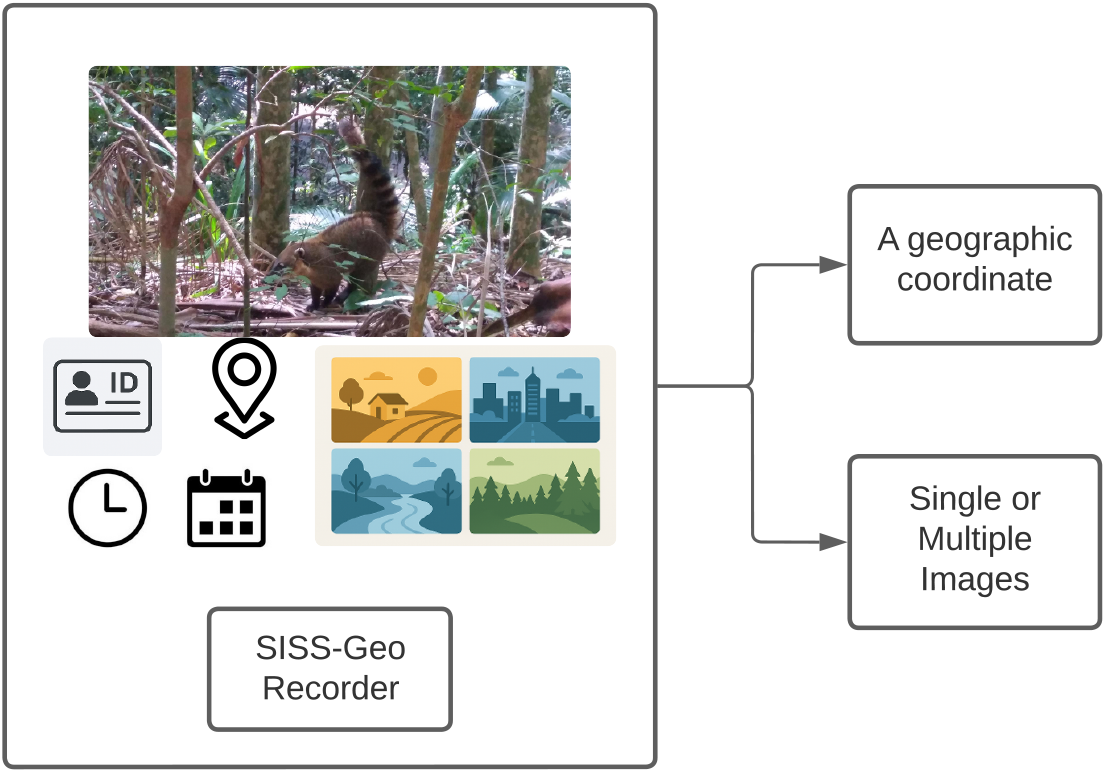
A detailed schematic representation of the SISS-Geo Recorder process, showing how the system captures single or multiple images and links each record to precise geographic coordinates for structured geospatial documentation and analysis.

Data augmentation is applied to promote a greater balance in the SISS-Geo dataset. The strategy aims to reduce disparities in the number of images per record and class, seeking a more uniform distribution of samples across the dataset. Although some classes naturally contain more records and more images per record—making perfect balance unattainable—the adopted approach strives to achieve the highest possible level of equilibrium. This balancing process is intended to improve the robustness and generalization capacity of both the image classification and species distribution models, ultimately supporting more effective model training.

Thus, for an effective comparison of MaxEnt, Bioclim, ResNet-50, ResNet-152, and the Integrated Model, the test set was separated. This set needs to contain each image associated with its geographic coordinates. Thus, to determine the test set data, ensuring that all data present an image and the geographic coordinates associated with the image, a random separation of 15% of the records is performed (percentage normally used in the literature) to be used for comparing the models. The training data are separated into data for the generation of image classification and species distribution models. Thus, we separated the data used in the MaxEnt training, which consists of the records of the classes of animals present in SISS-Geo. On the other hand, the images are used for training ResNet models.

After separating the data for training a MaxEnt and Bioclim models, the images were separated for application in ResNet-50 and ResNet-152. Images associated with records already selected for the test set are not considered in this separation, as well as the registers used in MaxEnt. Furthermore, to increase the set of available images, data augmentation procedures are applied to increase the availability of images that are used in training the image classification model. It is important to highlight that data augmentation operations were applied to the original images to balance the final data set for training the classification models. Thus, some classes suffered more increases than others. For example, the Monkey class did not go through the data augmentation process, as it is the class with the largest number of records available. Data augmentation is used to increase the number of images available for training deep networks, as better results are usually obtained when these models are trained with more images [31].

Since this is a highly unbalanced set, the focus of the data augmentation amplification was to balance the data set.

### 4.2 Integration of Models

Based on the Image Classification Model and the Species Distribution Model, integrated models are generated, which seek to combine the relative probabilities of each initial model. The integration follows the flow shown in Figure 9.

**FIGURE 9.**
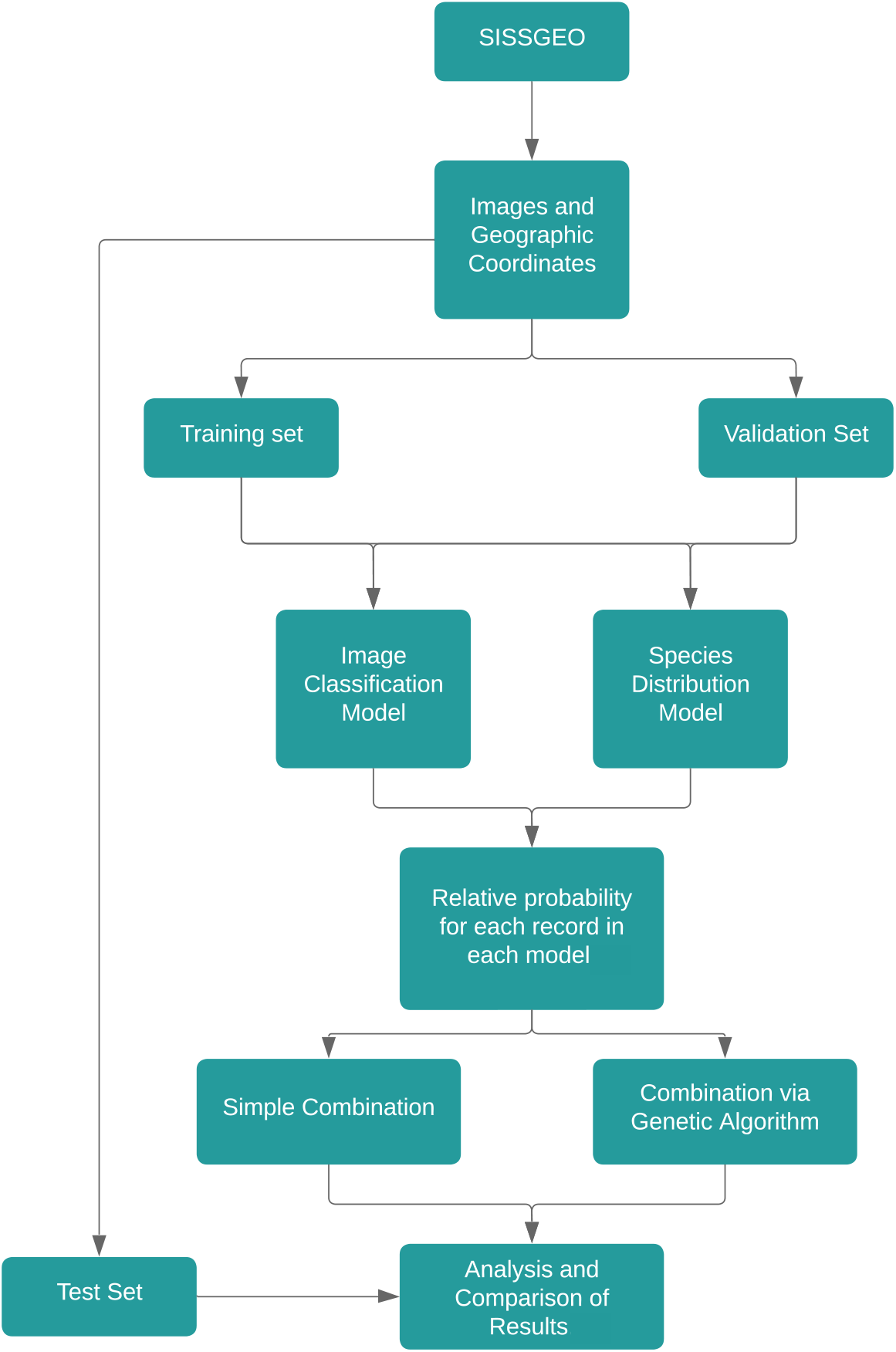
Integrated Model Workflow.

Two combinations of the models are performed. The first combination consists of using a genetic algorithm to obtain a single parameter *α*. While this first combination does not prioritize any specific classification model, the second combination tends to assign greater weight to the model that presents the best classification results. The combination is defined as:

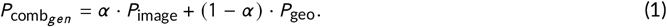

The parameter *α* is obtained through the genetic algorithm, which evaluates numerous variations of this parameter to find the value that best integrates the base models. The goal is to identify the weighting that produces the highest overall classification performance.

In addition to performing the combination with a single global parameter *α*, we also seek to combine the classification models by generating 15 parameters *α*_*i*_, where *i* = 1, …, 15. Each parameter corresponds to a specific class, allowing the combination to adapt to the behavior of the models for that class.

This approach is motivated by the observation that some species are better classified by the image-based model, while others show better results with the geographic model. Therefore, a class-dependent weighting may offer a more flexible and accurate combination.

The class-dependent combination is defined as:

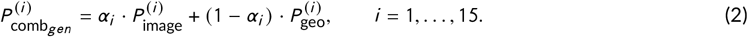

Here, 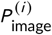 and 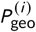 represent the predicted probabilities for class *i* obtained from the image-based and geographic-based models, respectively. The parameters *α*_*i*_ are optimized individually through the genetic algorithm, enabling a more refined and adaptive fusion strategy.

This class-dependent combination improves the overall flexibility of the model ensemble, as it allows the system to favor the model that performs better for each specific species.

To train the genetic algorithm, the validation set is used, where the relative probabilities of the image classification and species distribution models are present.

The proposed methodology integrates deep learning-based image classification with traditional species distribution modeling and evolutionary optimization to generate more robust predictive models. The workflow consists of three main stages: (i) image-based classification, (ii) species distribution modeling, and (iii) model combination through a genetic algorithm.

#### Stage 1: Image-based classification

The first stage relied on convolutional neural networks based on the ResNet architecture to classify images associated with different ecological classes. Two network depths were evaluated: ResNet-50 and ResNet-152. Both architectures were trained using a preprocessed image dataset, which included single-image and multi-image records. Each model produced a probability vector as output, representing the likelihood that a given record belonged to each target class.

#### Stage 2: Species distribution modeling

In parallel, two classical species distribution modeling (SDM) approaches were implemented: MaxEnt (Maximum Entropy) and BioClim. Unlike the image-based models, these methods use environmental and climatic predictors derived from each record’s georeferenced information. For every occurrence, both SDMs generated probability estimates that indicate the environmental suitability of each species or ecological class. These outputs were stored for subsequent integration with the image-based classification stage.

#### Stage 3: Model integration using a genetic algorithm

After obtaining probabilistic outputs from both modeling pathways (image-based and SDM-based), a genetic algorithm (GA) was employed to combine them into a unified predictive model. The GA optimized the weighting of the probabilistic contributions from each method by maximizing classification performance on a validation set. Through iterative evaluation, selection, recombination, and mutation, the GA identified the optimal combination strategy, yielding a final ensemble model that leverages complementary information from visual features and environmental suitability patterns.

In the hybrid modeling stage, two complementary combination strategies were explored to integrate the outputs of the image-based classifiers and the species distribution models. The first strategy consisted of a single-parameter model, in which the genetic algorithm optimized a global scalar weight that linearly combines the probabilities produced by both modeling pathways. This configuration imposes a uniform balance between visual and environmental information across all classes, offering a simple yet effective fusion mechanism. The second strategy adopted a more flexible multi-parameter formulation, in which the genetic algorithm optimized 15 independent weights—one for each target class. By allowing class-specific weighting, this approach captures variations in how strongly visual or environmental cues contribute to the classification of different taxa, leading to a more finely tuned ensemble. Together, these hybrid configurations enhance predictive accuracy by exploiting the complementary strengths of deep convolutional neural networks and classical species distribution models, resulting in a richer and more ecologically informed decision-making process.

## 5 RESULTS AND ANALYSIS

This section presents the results obtained with the methodology presented. Individual results are presented for each methodology applied. In addition, some observations for each method are presented. Finally, a comparison of the methods tested with the application on the set of images from SISS-Geo is shown.

The proposed analysis highlights that deep learning approaches (ResNet-50 and ResNet-152) achieve superior performance when multiple images per record are available. In contrast, single-record scenarios rely heavily on data augmentation to retain competitive results. Classical models such as MaxEnt and BioClim remain valuable as interpretable baselines, although they lack flexibility under complex environmental conditions. Finally, the genetic algorithm-based combinations produced the strongest outcomes overall, particularly with 15 environmental parameters, as they effectively integrate complementary strengths from both deep and classical learning paradigms.

### 5.1 Parameter Settings

All experiments were conducted under a unified experimental protocol in order to ensure reproducibility and a fair comparison among the evaluated models. The dataset was randomly divided at the record level, reserving 15% of the records for testing, 15% of the records for validation, and using the remaining 70% for training and validation, following common practice in the literature. Only records containing both image data and associated geographic coordinates were considered in the test set, which was kept fixed across all experiments. Random seeds were set to 42 in all implementations to reduce variability due to stochastic processes during training.

Species distribution models were trained exclusively using occurrence records associated with animal classes available in the SISS-Geo dataset. MaxEnt was configured using presence-only data and standard regularization parameters, with logistic output selected to generate probabilistic suitability estimates. Bioclim was trained using the same set of environmental variables to ensure methodological consistency and allow direct comparison between the two approaches.

For image classification, deep learning experiments were carried out using ResNet-50 and ResNet-152 architectures pre-trained on ImageNet. In both cases, the convolutional backbone was frozen during training, and only the final classification layers were optimized. The input images were resized to 224 × 224 pixels, and the number of output neurons in the final fully connected layer was set to 15, corresponding to the number of animal classes in the dataset. The models were trained using categorical cross-entropy loss and the Adam optimizer with a learning rate of 1 × 10^−4^. A batch size of 32 and a maximum of 60 training epochs were adopted for both architectures.

Early stopping was employed during training based on validation performance, with a patience of 10 epochs and automatic restoration of the best-performing model weights. Model evaluation was performed using the fixed test set, and classification performance was assessed through accuracy, confusion matrices, and per-class precision, recall, and F1-score. All experiments were implemented in Python, using TensorFlow/Keras for the ResNet-50 model and PyTorch for the ResNet-152 model, ensuring consistency with the original implementations provided.

The integration of model outputs was performed using two genetic algorithm (GA) configurations, designed to optimize the combination of probabilistic predictions generated by the image classification and species distribution models. In the first configuration, referred to as GA-1, a single global weight was optimized to balance the contribution of the two models across all classes, resulting in a simpler combination strategy with reduced dimensionality. In the second configuration, denoted as GA-15, a vector of 15 class-specific weights was optimized, allowing a finer and more flexible adjustment of the relative importance of each model for each animal class. In both cases, the genetic algorithm was initialized with a population of 100 individuals and evolved over 60 generations. Parent selection was performed using tournament selection, offspring were generated through arithmetic mean crossover, and mutation was applied with a high rate of 0.9 for GA-1 and a moderate rate of 0.5 for GA-15 to balance exploration and exploitation; however, each mutation introduced only a small perturbation (Gaussian noise with *σ* = 0.1, followed by clipping to [0, 1]), ensuring controlled and stable updates to the weights. An elitist strategy was adopted by preserving the top five individuals at each generation. Model fitness was evaluated using classification accuracy on the validation set, and the best solutions were subsequently assessed on the test set. To ensure robustness and reduce stochastic effects, all experiments were repeated using multiple random seeds ranging from 42 to 71.

### 5.2 Results of the Individual Models

The results indicate that ResNet-152 consistently outperforms ResNet-50 across all Top-*N* levels, demonstrating that increased network depth leads to superior discriminative capability. At the most restrictive level (Top-1), ResNet-152 achieves an accuracy of 64.13%, compared to 58.17% for ResNet-50, representing an improvement of approximately 6 percentage points.

As the value of *N* increases, both architectures show rapid performance gains; however, ResNet-152 maintains a systematic advantage. For instance, at Top-5 accuracy, ResNet-152 reaches 91.61%, whereas ResNet-50 attains 86.98%. At Top-10, the difference persists, with ResNet-152 achieving 99.01% compared to 96.58% for ResNet-50.

Furthermore, ResNet-152 converges to 100% accuracy at Top-13, while ResNet-50 only reaches full accuracy at Top-15. This earlier convergence highlights the greater robustness and representational capacity of the deeper architecture, particularly in applications where a limited number of candidate predictions is desired.

Figure 10 presents the confusion matrix of the proposed classification model considering 15 wildlife classes for ResNet-152. The results show a strong concentration of values along the main diagonal, indicating a high number of correct predictions for most categories. Nevertheless, some confusion can be observed between visually or taxonomically similar species, such as monkeys and marmosets, as well as among small mammals. These misclassifications highlight the intrinsic difficulty of distinguishing closely related classes and suggest that additional contextual or morphological features could further improve the model’s performance.

**FIGURE 10.**
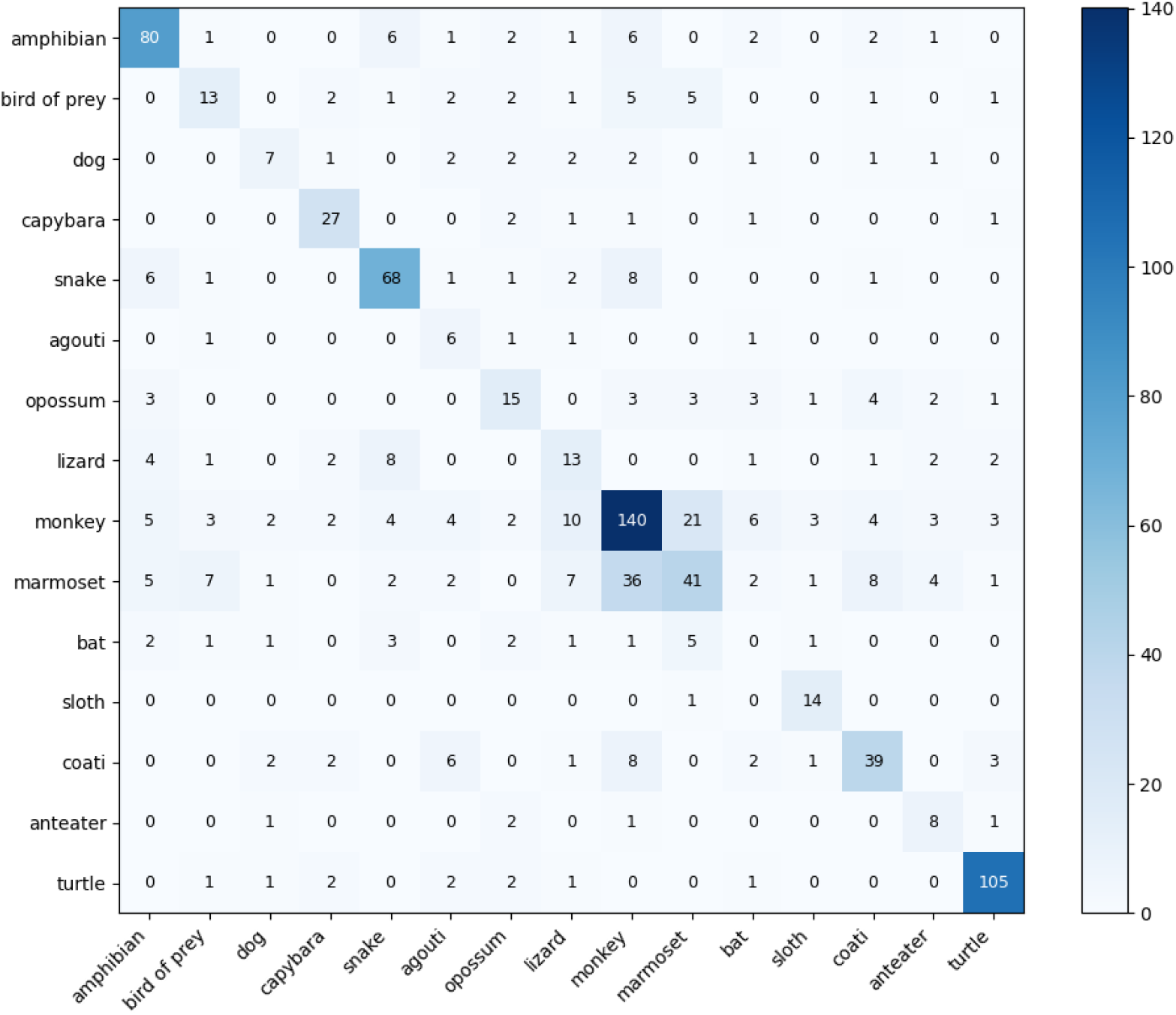
Confusion matrix obtained for the multiclass classification problem involving 15 wildlife categories. Rows represent the true classes and columns represent the predicted classes. Darker colors indicate higher frequencies, and the numerical values inside each cell correspond to the absolute number of samples.

When comparing the species distribution models, BioClim consistently outperforms MaxEnt across nearly all Top-*N* levels. At Top-1, BioClim achieves an accuracy of 31.76%, exceeding MaxEnt, which attains 27.79%. This result suggests that BioClim provides a more effective ranking of the most probable species under strict prediction constraints.

The performance gap becomes more pronounced at intermediate values of *N*. At Top-5, BioClim reaches 70.97%, outperforming MaxEnt by more than 12 percentage points (MaxEnt: 58.31%). A similar pattern is observed at Top-10, where BioClim achieves 93.55%, compared to 90.82% for MaxEnt.

Although both models converge to 100% accuracy at Top-15, BioClim consistently reaches higher accuracy levels earlier. This behavior indicates greater robustness and practical applicability in scenarios where the number of candidate species must be restricted. In contrast, MaxEnt exhibits a more gradual improvement, likely reflecting its sensitivity to pseudo-absence selection and the complexity of the environmental feature space.

### 5.3 Comparison between Deep Learning Models and Species Distribution Models

A clear performance gap is observed when comparing the deep learning models (ResNet-50 and ResNet-152) with the species distribution models (MaxEnt and BioClim). Across all Top-*N* values, the convolutional neural networks substantially outperform the SDMs, particularly at lower values of *N*, where prediction precision is most critical.

At Top-1, the best-performing CNN (ResNet-152) achieves 64.13%, more than double the accuracy of the best SDM (BioClim, 31.76%). This gap highlights the superior capacity of CNNs to extract discriminative features directly from image data, enabling more confident single-label predictions.

Even at intermediate levels, such as Top-5, CNN-based models exceed 85% accuracy, whereas SDMs remain below 71%. Although the performance difference narrows as *N* increases, CNNs consistently reach near-perfect accuracy with fewer candidate predictions. This earlier convergence suggests that deep learning models provide a more informative ranking of species probabilities.

Overall, while SDMs remain valuable for ecological interpretation and modeling species–environment relationships, CNN-based approaches demonstrate a markedly higher predictive power in image-based species identification tasks. These results suggest that deep learning models are better suited for applications requiring high classification accuracy, whereas SDMs may serve as complementary tools for ecological inference and spatial generalization.

Figure 11 presents a comparative analysis of the Top-*N* accuracy among the four evaluated methods: ResNet-50, ResNet-152, MaxEnt, and BioClim.

**FIGURE 11.**
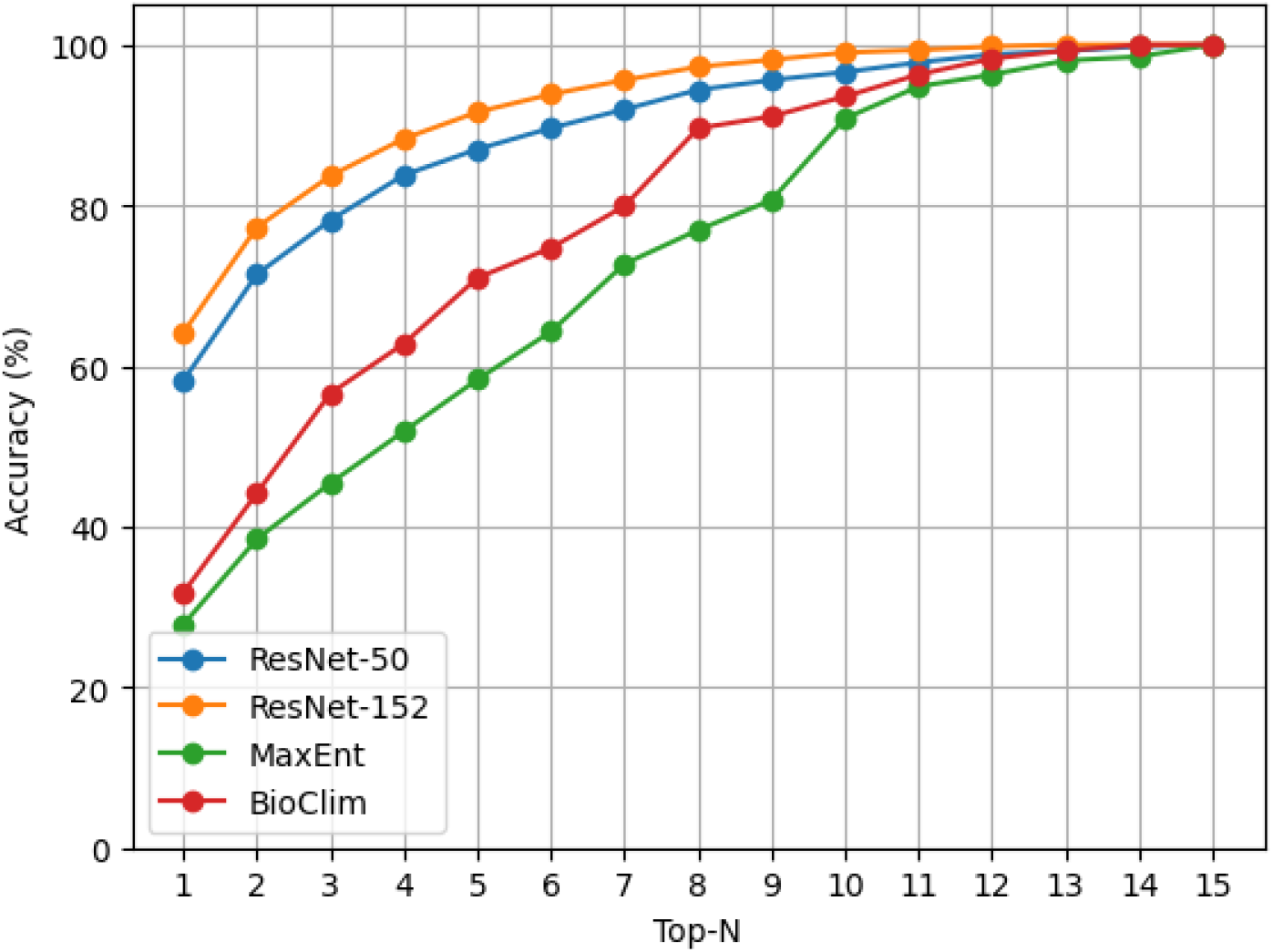
Accuracy Top-N - Single Models.

### 5.4 Results of the Integrated Models

The following subsection presents the results obtained from the integration of convolutional neural networks (CNNs) and species distribution models (SDMs) using a genetic algorithm (GA). The analysis is organized according to each model combination, allowing a detailed examination of their individual performance and integration behavior. For each case, both the single-weight and class-specific weighting strategies are considered, highlighting differences in accuracy, stability, and the relative contribution of visual and environmental information. In addition to the GA-based approaches, a simple baseline combination using equal weights (0.5 for each model) is also evaluated, providing a direct reference for comparison with the optimized integration strategies. This structured presentation provides a comprehensive understanding of how different model pairings and integration schemes influence the overall classification performance.

#### 5.4.1 ResNet-50 + MaxEnt

The integration of ResNet-50 with MaxEnt, using a single global weight optimized by a genetic algorithm (GA), achieved an average accuracy of 62.33%. The optimization process exhibited high stability across multiple runs with different random seeds, indicating consistent convergence.

The average weight assigned to the CNN was approximately 0.48, suggesting a slight predominance of the species distribution model (SDM) over visual features. This result indicates that environmental information plays a relevant complementary role in this configuration.

When class-specific weights were introduced, the performance improved to 63.95%. This gain highlights that different species benefit from distinct combinations of visual and environmental information.

#### 5.4.2 ResNet-50 + BioClim

The combination of ResNet-50 and BioClim achieved an average accuracy of 61.06% under the single-weight strategy. The average CNN weight was approximately 0.59, indicating a stronger reliance on visual features compared to the climatic model.

With class-specific weighting, accuracy increased to 61.94%, although the improvement was more modest compared to the MaxEnt-based integration. This suggests that, in this configuration, the CNN already dominates the classification process for most classes.

#### 5.4.3 ResNet-152 + MaxEnt

The integration involving the deeper ResNet-152 architecture led to substantial improvements. Under the single-weight strategy, the ResNet-152 + MaxEnt combination achieved an average accuracy of 68.63%.

Additionally, a simple ensemble using equal weights for both models (0.5 for the CNN and 0.5 for MaxEnt) was evaluated as a baseline integration strategy. This configuration achieved a Top-1 accuracy of 58.61%, reaching 77.48% at Top-5 and 95.03% at Top-10. These results demonstrate that even a non-optimized combination is capable of effectively leveraging complementary visual and environmental information.

This highlights the importance of ecological constraints in the classification task, even in the presence of a highly expressive CNN architecture.

When class-specific weights were applied, performance increased significantly to 70.62%, representing the best result among all evaluated configurations. This improvement reinforces the importance of class-level adaptation in hybrid model integration.

#### 5.4.4 ResNet-152 + BioClim

The ResNet-152 + BioClim combination achieved the best performance under the single-weight strategy, with an average accuracy of 69.28%.

A baseline ensemble using equal weights (0.5 for each model) was also analyzed. This approach achieved a Top-1 accuracy of 54.30%, while reaching 98.23% at Top-5 and 99.56% at Top-10. The results indicate that BioClim contributes useful complementary environmental information, particularly for broader ranking thresholds, even without optimization.

With class-specific weights, the accuracy reached 68.58%, slightly lower than the MaxEnt-based configuration. This suggests that, although BioClim performs well under global integration, its class-level discriminative power may be more limited.

### 5.5 Comparative Analysis of the Results

The results consistently demonstrate that integrating convolutional neural networks (CNNs) with species distribution models (SDMs) outperforms standalone approaches. While SDMs alone yield relatively low accuracy (MaxEnt: 27.79%, BioClim: 31.76%), CNNs achieve significantly better performance (ResNet-50: 58.17%, ResNet-152: 64.13%). Furthermore, even a simple ensemble strategy based on equal weights (0.5 for the CNN and 0.5 for the SDM) produced competitive results, achieving 58.61% Top-1 accuracy for the ResNet-152 + MaxEnt combination and 54.30% for ResNet-152 + BioClim, while substantially improving Top-*k* performance at broader ranking thresholds.

The use of a genetic algorithm to combine models leads to additional improvements across all configurations. Even when optimizing a single global weight, the hybrid models show consistent gains, indicating that visual and environmental information are complementary rather than redundant.

The most significant improvements are observed when class-specific weights are employed. This strategy allows the model to capture inter-class heterogeneity by dynamically adjusting the contribution of each component.

Figure 12 summarizes the Top-1 accuracy results across all evaluated models and integration strategies. The figure reveals a clear performance hierarchy. Standalone SDMs present the lowest accuracy, followed by CNN-only models, while all integrated approaches outperform their individual counterparts. Notably, the ResNet-152 combined with MaxEnt using class-specific weights achieves the highest Top-1 accuracy, confirming the effectiveness of both deeper architectures and adaptive integration.

**FIGURE 12.**
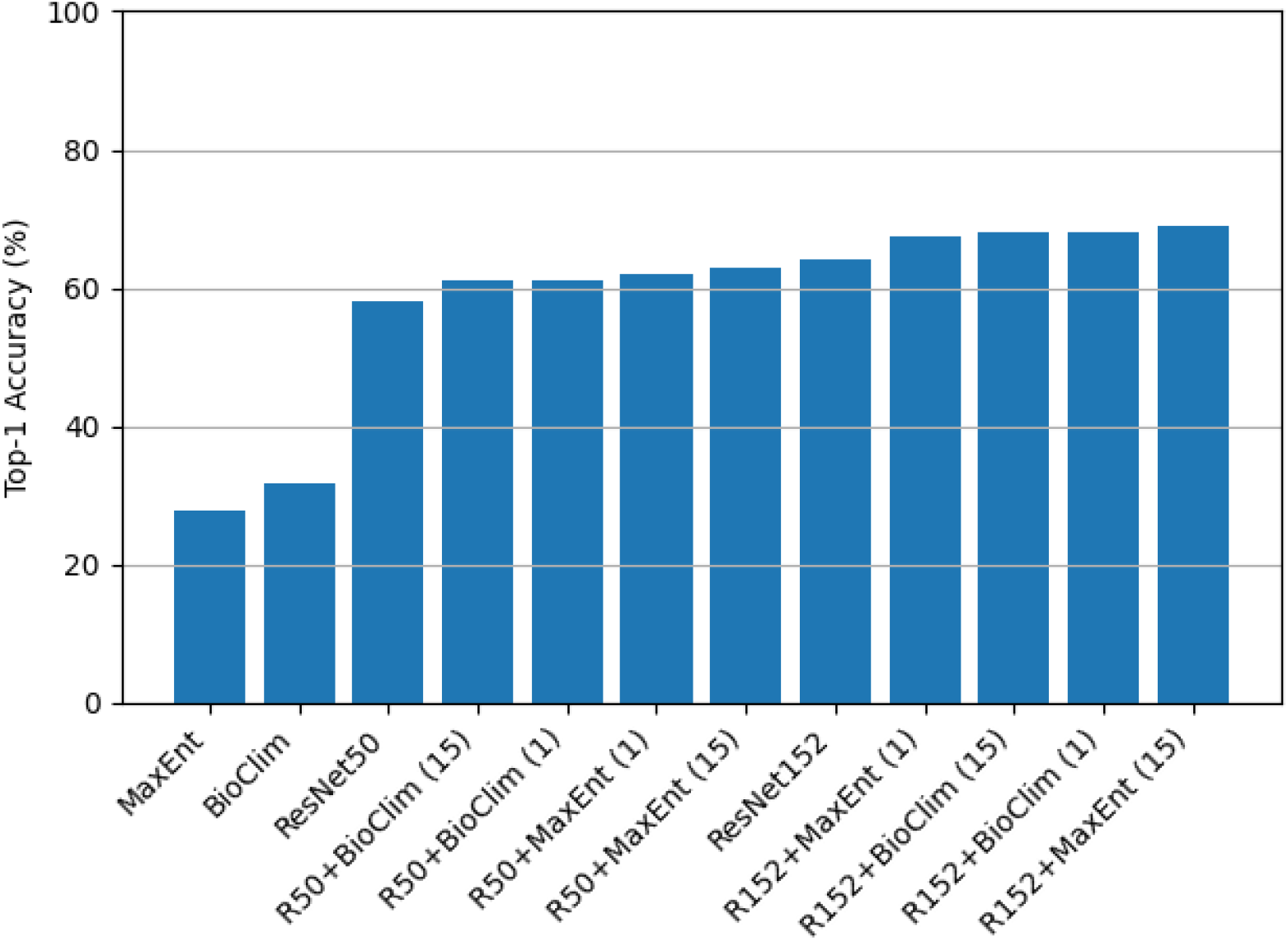
Top-1 Accuracy Comparison Across Models and Integration Strategies.

Network depth plays a crucial role in this process. The deeper ResNet-152 architecture consistently outperforms ResNet-50 and shows greater capacity to exploit complementary information from SDMs.

An important advantage of the GA-based integration is its interpretability. The learned weights provide meaningful insights into the relative importance of visual and environmental information across species.

The Top-*N* accuracy analysis further supports these findings. Integrated models outperform both CNN-only and SDM-only approaches at Top-1, while maintaining competitive performance as *N* increases.

Figure 13 presents the Top-*N* accuracy curves. The integrated models show clear improvements in the most critical positions (low *N*), indicating better ranking of the correct class. As *N* increases, all models converge toward high accuracy levels; however, the integrated approaches maintain a consistent advantage, demonstrating more robust ranking behavior.

**FIGURE 13.**
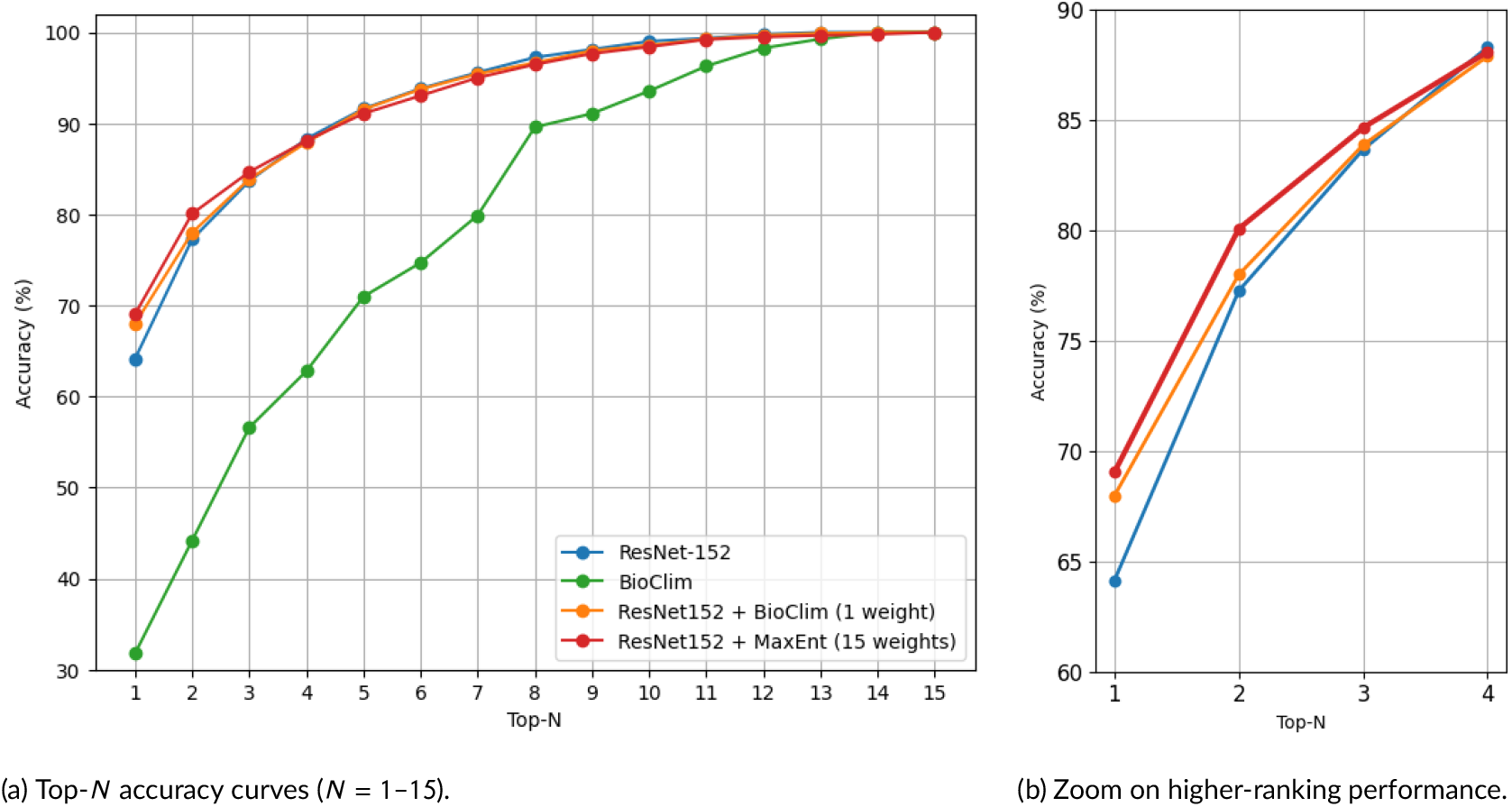
Comparison of Top-*N* accuracy across models. The integrated approaches improve Top-1 performance and maintain superior ranking quality, converging to high accuracy as *N* increases.

In summary, the results show that model integration consistently improves performance, deeper CNNs enhance the gains, and class-specific weighting is essential to capture inter-species variability. The genetic algorithm provides a robust, interpretable, and effective framework for combining heterogeneous models, reinforcing the potential of hybrid approaches that bridge computer vision and ecological modeling.

## 6 PERSPECTIVES AND APPLICATIONS

The results obtained in this study demonstrate the benefits of integrating image-based deep learning models with species distribution models for automated wildlife classification. When evaluated independently, image classification models achieved substantially higher performance than species distribution models, with ResNet-152 reaching a Top-1 accuracy of 64.13

The methodology proposed in this work opens several opportunities for practical applications and future developments in biodiversity monitoring, wildlife health surveillance, and ecological research. By combining visual information extracted from images with ecological information derived from species distribution models, the framework provides a robust mechanism for supporting automated species identification in large-scale monitoring systems.

One immediate application is the integration of the proposed approach into citizen-science and environmental surveillance platforms such as SISS-Geo. Automated identification tools can assist specialists in validating incoming records, prioritizing uncertain observations for expert review, and accelerating the processing of large volumes of wildlife data. Such capabilities are particularly relevant in countries with high biodiversity and limited taxonomic expertise available for manual validation.

From an ecological perspective, the integration of image classification and species distribution modeling can contribute to biodiversity conservation by improving the quality and reliability of species occurrence records. More accurate species identification supports ecological monitoring programs, habitat management initiatives, invasive species detection, and conservation planning efforts. Furthermore, the explicit incorporation of environmental information allows the system to consider ecological constraints that may not be evident from visual information alone.

The proposed framework also has important implications for wildlife health surveillance and zoonotic disease monitoring. Many species recorded in SISS-Geo are relevant to epidemiological surveillance due to their potential roles as disease reservoirs, hosts, or vectors. Improved species identification can therefore enhance the quality of information available to environmental and public health agencies, supporting early warning systems and risk assessment strategies associated with emerging infectious diseases.

Beyond the specific application explored in this study, the methodology can be extended to other multimodal ecological problems involving heterogeneous sources of information. Future developments may include the incorporation of higher-resolution environmental variables, additional contextual information available in SISS-Geo records, and advanced multimodal architectures capable of learning joint representations of visual, spatial, climatic, and ecological data. Adaptive weighting strategies could also be extended to dynamically adjust model contributions according to prediction confidence, environmental uncertainty, or species-specific characteristics.

More broadly, this work demonstrates the potential of integrating artificial intelligence techniques with ecological modeling to support evidence-based environmental management. As biodiversity monitoring programs continue to expand and generate increasingly large datasets, multimodal approaches such as the one proposed here may become valuable tools for transforming raw ecological observations into actionable information for conservation, wildlife management, public health, and sustainable environmental governance.

## Abbreviations

SDM: species distribution model
CNN: convolutional neural network
GA: genetic algorithm

## Acknowledgements

The authors thank the financial support provided by the Brazilian agency Conselho Nacional de Desenvolvimento Científico e Tecnológico (CNPq) (grants 445493/2023-2 and 313452/2025-3), and FAPEMIG (grants APQ-01832-22 and APQ-03313-22). The Article Processing Charge (APC) for the publication of this research was funded by the Coordination for the Improvement of Higher Education Personnel - CAPES (identifier ROR: 00×0ma614). For open access purposes, the authors have assigned the Creative Commons CC BY license to any accepted version of the article. We also acknowledge the support provided by UFJF and Fiocruz. This work has been supported by High-Speed Integrated Research Network (RePesq).

## DATA AVAILABILITY STATEMENT

The source code, scripts, and implementation details used in this study are publicly available in the following GitHub repository: https://github.com/Mateusbo247/species-classification.git. The repository includes the experimental pipeline, model configurations, and supporting resources necessary to reproduce the reported results.

The occurrence records and associated species distribution maps used in this study are publicly available through the Brazilian Biodiversity Information System (SISSGEO) at https://sissgeo.lncc.br/mapaRegistrosInicial.xhtml. These data can be accessed and visualized through the platform’s public interface.

## Footnotes

1 https://sissgeo.lncc.br/apresentacao.xhtml

2 https://www.worldclim.org/data/bioclim.html

3 https://csiropedia.csiro.au/bioclim/

4 https://cran.r-project.org/web/packages/dismo/

